# Microtubule Disruption does not Impair Learning in the Unicellular Organism Paramecium: Implications for Information Processing in Microtubules

**DOI:** 10.1101/712794

**Authors:** A. Alipour, G. Hatam, H. Seradj

## Abstract

Information processing in microtubules is an open question that has not been properly addressed yet. It was suggested that microtubules could store and process information in the nervous system or even support consciousness. The unicellular organism, *Paramecium caudatum*, that has a microtubular structure but does not have a neuron or neural network, shows intelligent behaviors such as associative learning. This may suggest that the microtubules are involved in intelligent behavior, information storage or information processing in paramecium. To test this hypothesis, we have utilized a paramecium learning task in which the organism associates brightness in its swimming medium with attractive cathodal shocks to study the role of microtubules in paramecium learning. We used an antimicrotubular agent (parbendazole) and disrupted microtubular dynamics in paramecium to see if microtubules are an integral part of information storage and processing in paramecium’s learning process. We observed that a partial allosteric modulator of GABA (midazolam) could disrupt the learning process in paramecium, but the antimicrotubular agent could not. Therefore, our results suggest that microtubules are probably not vital for the learning behavior in *P. caudatum*. Consequently, our results call for a further revisitation of the microtubular information processing hypothesis.

## I. INTRODUCTION

Herbert Froehlich was one of the first to suggest the feasibility of long life macroscopic coherence (classical and quantum) in “ordered” biological structures such as microtubules at room temperature (1,2). Furthermore, some researchers argued that coherence between microtubules in the nervous system could potentially provide an explanation for consciousness (3,4). While the issue of quantum effects in the human brain is a controversial topic of science (5,6) and it is likely that decoherence does not let quantum states to survive long enough to be effective in cognitive processing (7–9), the possibility of information processing in microtubules is still an open question.

In a series of studies by G Albrecht-Buehler, he has demonstrated that cell intelligence in fibroblast cells is due to some cytoskeletal structures in the cell, and living cells possess a spatial orientation mechanism located in the centriole (i.e. a microtubular structure), and electromagnetic signals are the triggers for the cells repositioning (10–12). Meanwhile, it is still controversial how the reception of electromagnetic radiation is accomplished by the centriole.

Moreover, some argued that learning in unicellular organisms such as paramecium might support the hypothesis that sub-cellular structures (such as microtubules) could support intelligent behavior (13). In a similar vein, it was suggested that microtubules do not only process information, but they are also responsible for memory storage and learning. A modelling study showed that the structure of Ca2+/calmodulin-dependent protein kinase II (CaM kinase II or CaMKII which is essential for memory formation in neurons) fits into the phosphorylation sites on tubulins of microtubules (14). This strengthens the possibility that tubulin phosphorylation in microtubules may encode information as it was suggested elsewhere(15).

Here, we sought to test the hypothesis that microtubules are involved in memory encoding and information storage in a unicellular organism that exhibits learning. We disrupted the microtubular dynamics in the organism to see if microtubules are involved in information storage in paramecium. Our results suggested that while disrupting GABA receptor dynamics will impair paramecium learning, disruption of microtubular dynamics does not impair the learning behavior.

## II. METHODS

### A. Experimental Setup

To investigate the role of microtubules in paramecium learning, we used a previously developed behavioral learning paradigm (16,17) and combined it with a pharmacological manipulation approach (18). Accordingly, we tested the effect of two different drugs on paramecium’s learning capability to investigate its learning mechanisms. Specifically, to investigate the role of microtubules in paramecium learning we disrupted the dynamics of microtubules in paramecium using parbendazole. Moreover, we investigated the role of GABA receptors in paramecium learning through the application of Midazolam, a benzodiazepine and a GABA partial allosteric modulator (GABA PAM), to find other possible mechanisms for this process.

### B. *Paramecium caudatum* specimens

The experimental procedure of this experiment was reviewed by the ethics committee of the shiraz university of medical sciences (designated code: IR.SUMS.REC.1394.S999, available at http://research.sums.ac.ir/fa/ethicrc/EthicsCodes.html). Local samples of Khoshk River in the city of Shiraz, Iran were gathered and incubated in hey infusion as the nutritious culture medium for paramecia with a volumetric ratio of 1 to 10 (i.e. 10 ml of a specimen with 100 ml hey infusion). After 3 days, the specimens were checked under 40X magnification for the presence of paramecia and isolated for further evaluation. Paramecium caudatum was identified based on its unique morphological features, i.e. the large size (∼300 micrometres) and the presence of only one micronucleus beside the large macronucleus. Afterwards, isolated paramecia were added into a new culture medium (20 ml of hey infusion) and incubated in culture flasks in room temperature. Cultures were passaged every 3 days by replacing 1/5 of the previous culture with fresh culture medium. In this study, the culture medium was not axenic.

### C. Learning behavior in *P. caudatum*

The methodology of Armus et al. (19) was used to investigate the learning behavior of P. caudatum. A U-shaped plastic trough (20mm length, 5mm width, and 5mm depth) was filled with the original culture medium that the test paramecium was isolated from (which will be called swimming medium from now on). In order to exclude other microorganisms/impurities from the culture medium in the trough, the culture medium had been filtered through a 0.22-micrometer filter. Copper-ended cathode and anode wires were placed at the center of the side walls at two ends of the trough. A length of 3-millimeter copper end of the wire was exposed to culture medium at each end. The trough was divided into two dark and light sides using a dark transparent sheet that was placed underneath the plastic trough. Light intensity was set to 80530 lux for the bright side and 33530 lux for the dark side of the trough (See figure 1 for more detail).105 Paramecia (P. caudatum) were divided into four groups of experimental, control, parbendazole and midazolam treated.

**FIG. 1.**
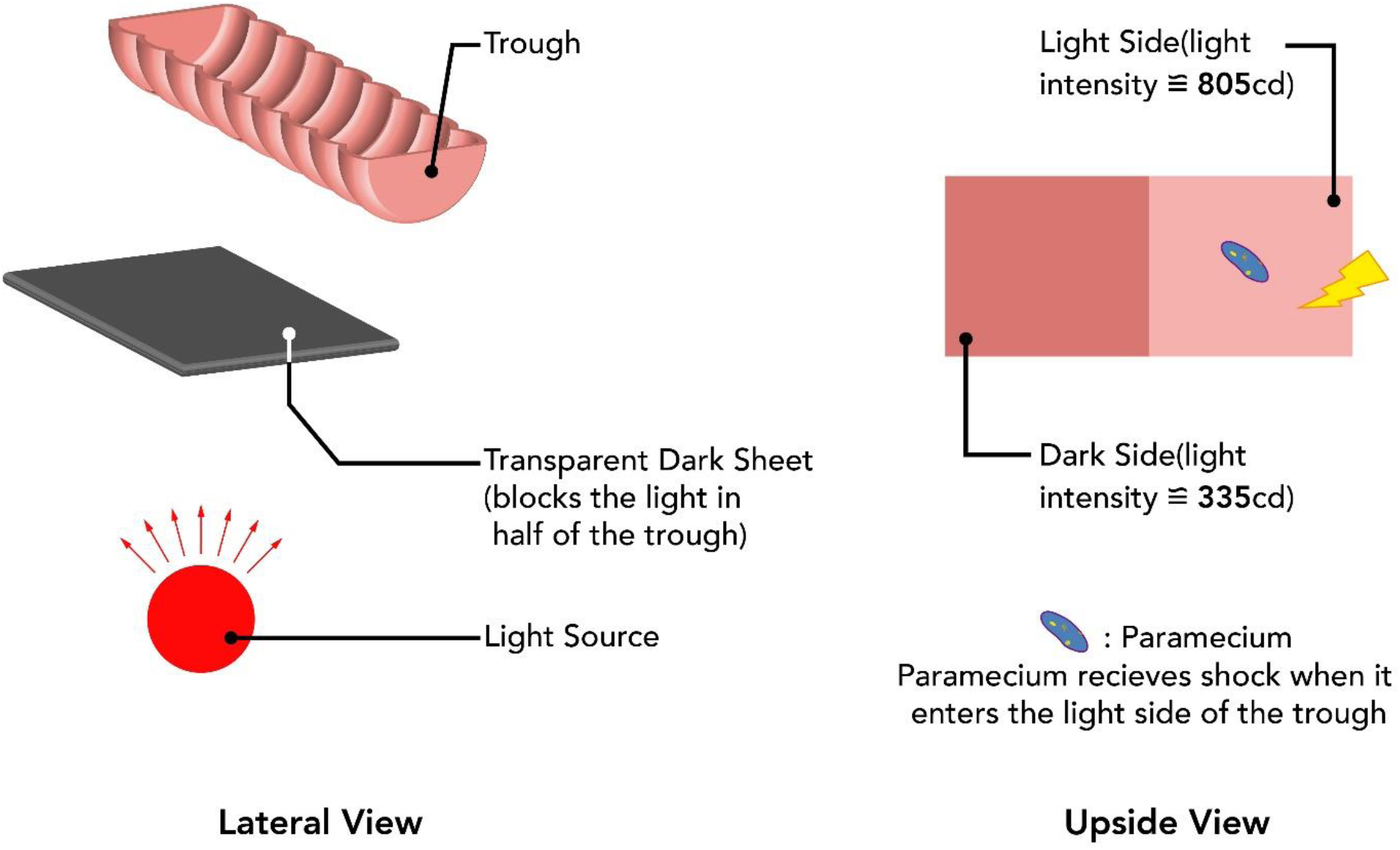
A schematic representation of the experimental setup.

For the experiment, each paramecium in experimental or control group underwent ten 90-second trials, 7 training trials, and 3 test trials without any inter-trial time intervals. The swimming medium between trials was not changed (to prevent possible confounding factors such as changing paramecium’s environmental cues, lighting conditions, interference with free exploration of the paramecium, etc.) and individual paramecia were watched uninterrupted during the whole training and test trials. To place the paramecium into the swimming medium, a few drops of culture flask containing paramecium were placed on a microscope with 40X magnification and single paramecia were sucked into a mouth pipet tipped with a capillary tube (diameter of ∼500 micrometers). Subsequently, paramecia were injected to the swimming medium and observed under 10X magnification of a stereo microscope. In training trials of the experimental group (the first 7 trials), the experimenter manually started the shocks when the paramecium entered the cathodal half of the trough (weather it was the dark or bright side) and shocks were turned off when the organism left that half. In the test trials, the total time that paramecium spent in the light side of the trough was recorded. In the control group, paramecia never received any shock and the time spent in the dark and light sides of the trough in the test trials was recorded. Parbendazole treated group underwent the same procedure as the experimental group with the difference that the parbendazole group was treated with 100 micromolar solution of parbendazole for 24 hours before starting the experiment. For the midazolam group, paramecia received the same treatment as the experimental group with the difference that the swimming medium during the experiment contained 30 micrograms/ml of midazolam.

### D. Parbendazole

Parbendazole is a member of benzimidazole anthelmintic drug family that has been used to treat parasitic infections in veterinary settings (20). Even though parbendazole has been reported to interfere with cell metabolism as an inhibitor of glucose uptake and fumarate reductase and to disrupt secretory processes of acetylcholine (20,21), its main mechanism of action is to inhibit microtubule polymerization inside the cytoplasm(22). Consequently, due to the depolymerization of microtubules during the so-called dynamic instability of microtubules, parbendazole effectively makes microtubular depolymerization irreversible. This results in disintegration of microtubular networks inside the cytoplasm overtime. For example, application of 20 micromolar parbendazole in Vero cells for 21 hours only leaves centrioles intact and causes an almost complete disappearance of microtubules(22).

More importantly, parbendazole is an ideal agent to study the role of microtubular dynamics in paramecium since it does not cause cell death even at high concentrations (23). More specifically, Pape et al. tested the effect of a wide variety of antimicrotubular agents on paramecium’s growth and found that although parbendazole can effectively inhibit cell growth in paramecium, it does not show cytotoxic effects (23). Therefore, we have chosen parbendazole to study the relationship between microtubular dynamics disruption and paramecium learning while avoiding possible cytotoxic effects of antimicrotubular agents. To ensure that the parbendazole affects microtubules in paramecium, we chose a dose that was 2 orders of magnitude higher than the effective dose reported by Pape et al. (23) and 10 times bigger than what has been reported to virtually inhibit all microtubular assembly *in vitro* (22).

### 1. Parbendazole treatment

We incubated paramecium in a culture solution with 100 micromolar concentration of parbendazole for 24 hours prior to the experimental test for the parbendazole group. This time interval was selected based on Pape et al. report (23) indicating the apparent presence of parbendazole’s effect after 24 hours. Parbendazole disrupts microtubular dynamics of the paramecium and it should negatively affect the learning behavior of paramecium if microtubules are involved in paramecium’s learning. Parbendazole was acquired from Sigma-Aldrich, USA.

### E. Midazolam

Midazolam is a benzodiazepine drug that is widely used in medical procedures such as anesthesia, sedation, and amnesia (24). Midazolam is a partial allosteric modulator of GABAA receptors. While it does not cause direct opening of the GABAA receptors, it increases the frequency of opening in these receptors upon agonist binding (25). This results in an increased inflow of Cl− and hyperpolarization of target cells. Inspired by the amnesic effects of the midazolam in human subjects, we sought to test the idea that similar effects can be observed in paramecium due to evolutionary preserved pathways for memory formation across different species. To make the midazolam dosage comparable to parbendazole, we chose a dosage of midazolam that was 2 orders of magnitude higher that the previously reported plasma concentration of midazolam required for maintenance of anesthesia in human subjects (i.e. 30 micrograms/ml in the experiment as compared to reported values of 259 to 353 ng/ml in human subjects (26)).

### 1. Midazolam treatment

A filtered cultured medium with 30 micrograms/ml (≈90 nM) concentration of midazolam was prepared and used as the swimming medium for P. caudatum in learning trials of the midazolam group. To test the role of GABA receptors in learning behavior of paramecium, we used Midazolam as it disrupts the normal functioning of GABA receptors. Midazolam was acquired from Exir pharmaceutical company, Iran.

### F. Effect of parbendazole on paramecium’s growth

To ensure that parbendazole is affecting P. caudatum’s microtubular dynamics, we measured the growth of paramecia after parbendazole administration as a proxy for parbendazole’s effectiveness since population growth has a direct relationship with healthy microtubular dynamics while other known pharmacological effects of parbendazole do not directly affect the population growth and cell division. A sample population of paramecia was taken and divided into two groups of control and parbendazole (with 100 micromolar concentration). Then, the population density of paramecia was counted in each of the samples before, and 24 hours after parbendazole administration to the parbendazole group. To estimate the population density in each sample, we collected 28 samples (20 microliters each) from each culture mediums and counted the number of paramecia in each sample using an optical microscope. As stated earlier, the population density was recorded both at the parbendazole inoculation time and 24 hours after inoculation.

This procedure was performed in both logarithmic and stationary growth phases of paramecia. As reported before, the effect of antimicrotubular agents on logarithmic growth phase can be studied by their administration 24 hours after a new passage (23). For the stationary phase, we evaluated the effect of parbendazole 72 hours after a new passage.

### G. Effect of parbendazole on *P. caudatum* swimming speed

The swimming speed of the P. caudatum was recorded and measured using a microscope mounted camera before and 24 hours after parbendazole inoculation. Specifically, a sub-population of paramecia were randomly sampled, and the movement of paramecia was filmed for 10 minutes using the camera that was mounted on top of an eyepiece with a scale bar. Subsequently, movement speed (micrometers/second) of randomly selected paramecia was calculated by measuring the amount of movement of each paramecium under the microscope and a mean speed was calculated based on population average (n=10 for both groups).

### H. Electrical shock device

A microcontroller was used to adjust the shocks to the culture medium (ATMEGA 16 AVR controller). The circuit was designed to deliver cathodal shocks of 5 volts (60-millisecond shocks with 500-millisecond no-shock intervals) as soon as the experimenter pressed the bottom (which happened when the paramecium entered the bright side of the trough, see learning behavior in paramecium section for more information).

### I. Statistical analysis

We used one-way ANOVA (Tukey HSD for post hoc) in order to analyze the time spent in the bright side of the trough. This procedure was used to determine the time differences between four groups of control, experimental, parbendazole and midazolam. SPSS software (version 16.0) was used for statistical analysis. The effect of parbendazole and midazolam on the population and swimming speed of the P. caudatum were evaluated using the independent t-test in the same program.

## III. RESULTS

### Differences in behavioral profiles

Out of the 270 seconds time in the test period, P. caudatum learning group and parbendazole treated group spent 152±12.7 and 137±12.5 seconds in the bright side, respectively. This turned out to be significantly longer when compared to 105±8.1 seconds in the control group (one-way ANOVA, Tukey HSD test, P<0.01 and P<0.05 for learning group and parbendazole group, respectively. See figure 2 for more detail). For midazolam group, however, we did not observe a significant increase in the time spent inside the bright side (126±8.7 seconds, P=0.31). All numbers are reported in the mean±SEM format.

**FIG. 2.**
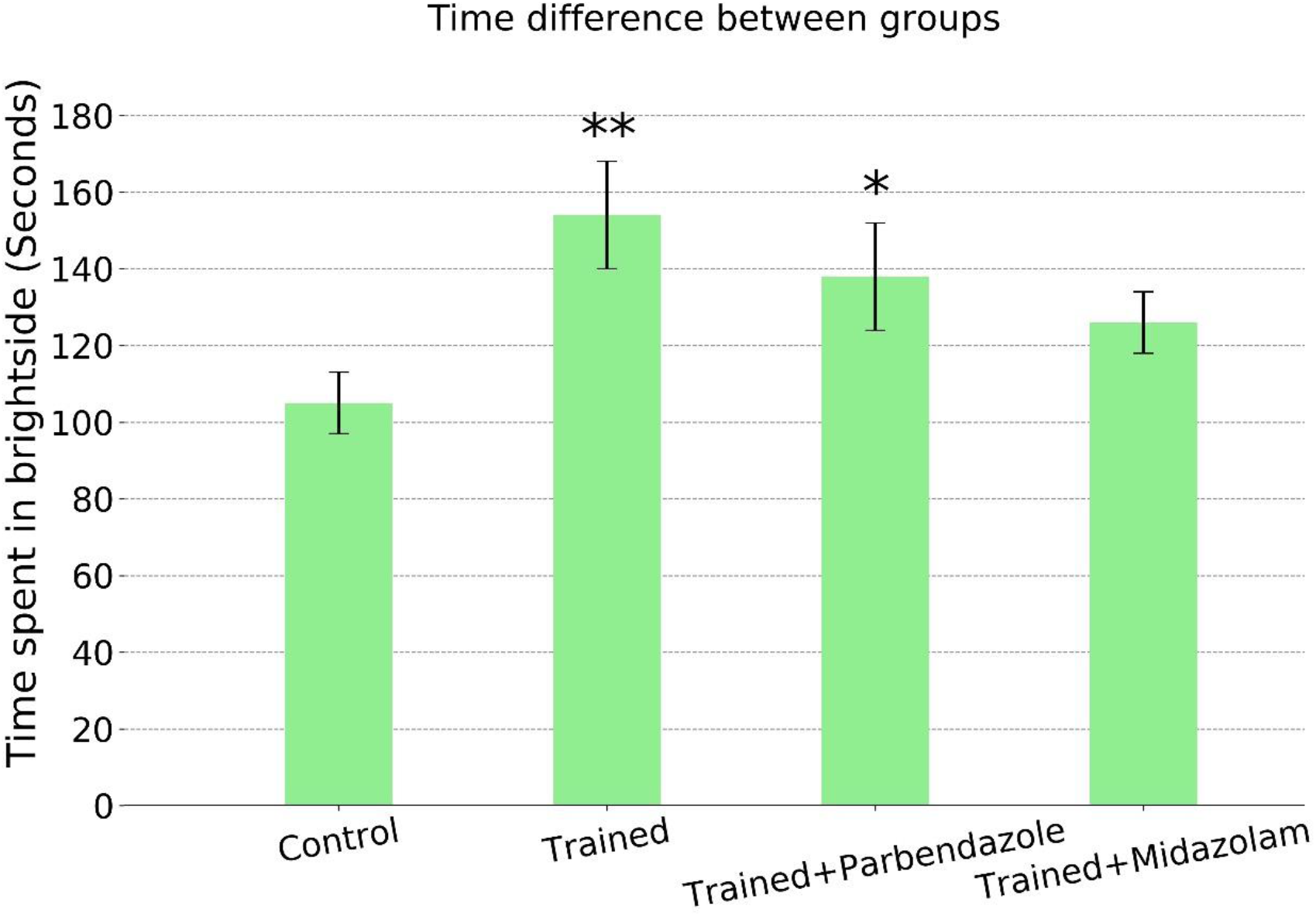
Time difference between groups: Control (n=30), trained without any drug (n=23), trained-parbendazole treated (n=26), and trained-midazolam treated group (n=26). While there was a significant difference between the time of the trained/parbendazole groups and control (one-way ANOVA, Tukey HSD test, P<0.01 and P<0.05, respectively), midazolam treated group did not show a significant time difference compared to the control group. Error bars are ±SEM.

### Drug effectiveness analysis for parbendazole

Drug effectiveness analysis was performed in both logarithmic and stationary phases. For the stationary phase, while at the time of the inoculation the parbendazole treated group and control group had the same population density, after 24 hours parbendazole treated paramecia had a significantly lower population density comparing with the control group (figure 3). In the logarithmic growth phase, control population showed a significant increase in the number of cells while the parbendazole treated group demonstrated an inhibition of population growth (figure 4). Note that since each single paramecium is being tested individually, population density does not affect learning behavior of paramecium.

**Figure 3.**
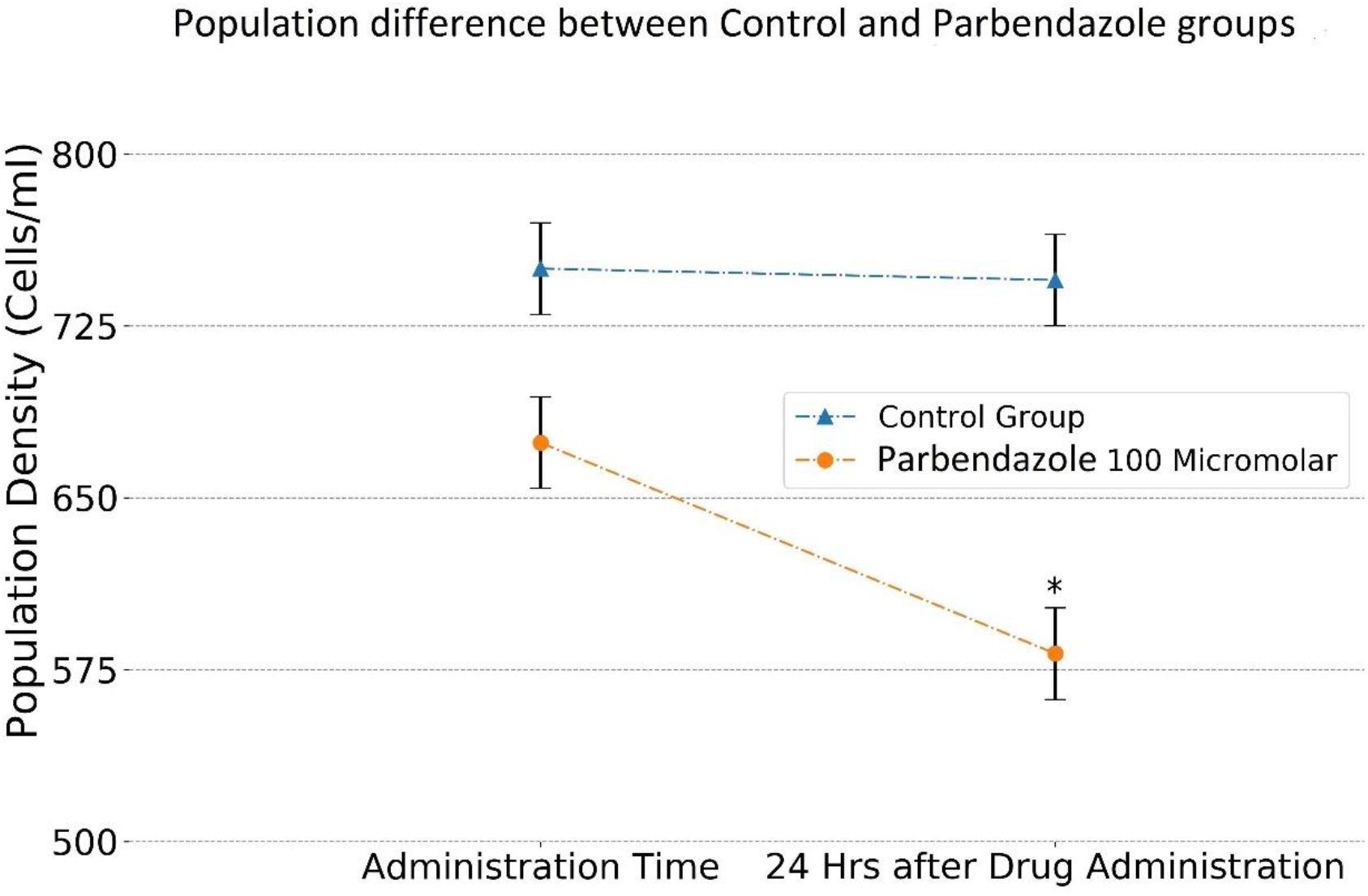
The population density of paramecium. Population density of *P. caudatum* before and 24 hours after the administration of parbendazole (dash-dotted line) compared to the control group (dotted line). While there was not any significant difference between two groups at the administration time, t-test indicated a significant difference after 24 hours between two groups in the stationary phase (p<0.05, n=28), error bars are ±SEM. Note that the reduction in population density is due to stationary phase of the growth i.e. while population stays the same in control group due to equal rates of division and cell death, an inhibition of cell division in the parbendazole group causes a net reduction of population density decline. Figure 4 shows that this effect is unlikely to be due to cytotoxicity.

**FIG. 4.**
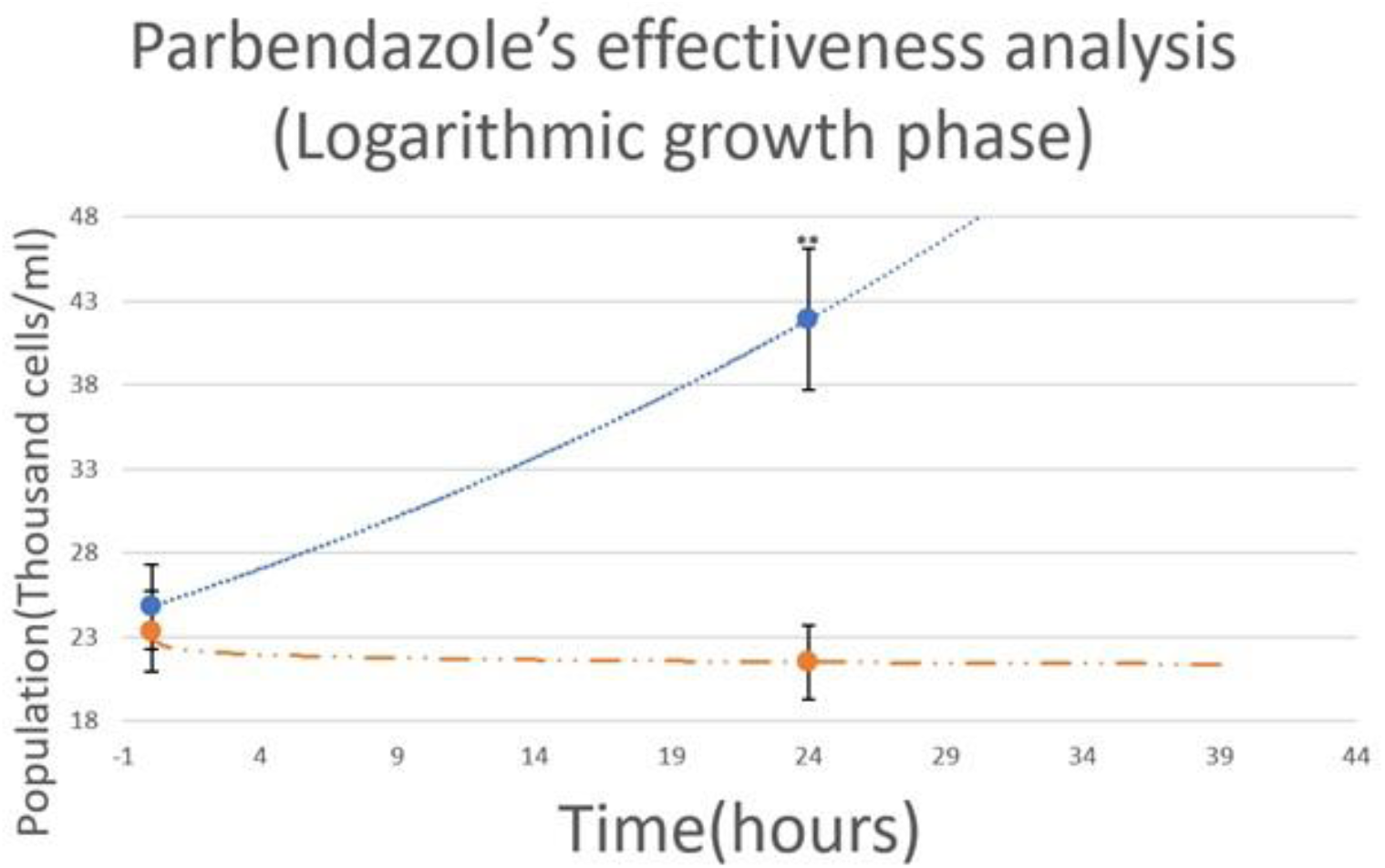
The population density of paramecium. In the logarithmic growth phase, the control group (dotted line) shows a logarithmic growth while the parbendazole group (dash-dotted line) demonstrates a growth inhibition due to the parbendazole administration after 24 hours (*p*<0.01, n=28), error bars are ±SEM. Lack of population density decline in logarithmic phase suggests that parbendazole was not cytotoxic at this concentration.

### Paramecium swimming speed analysis for parbendazole group

The independent *t*-test indicated that the swimming speed difference of P. caudatum before and after parbendazole treatment was not significant (1031±131 and 969±107 micrometers per second, respectively. See figure 5 for more detail).

**Figure 5.**
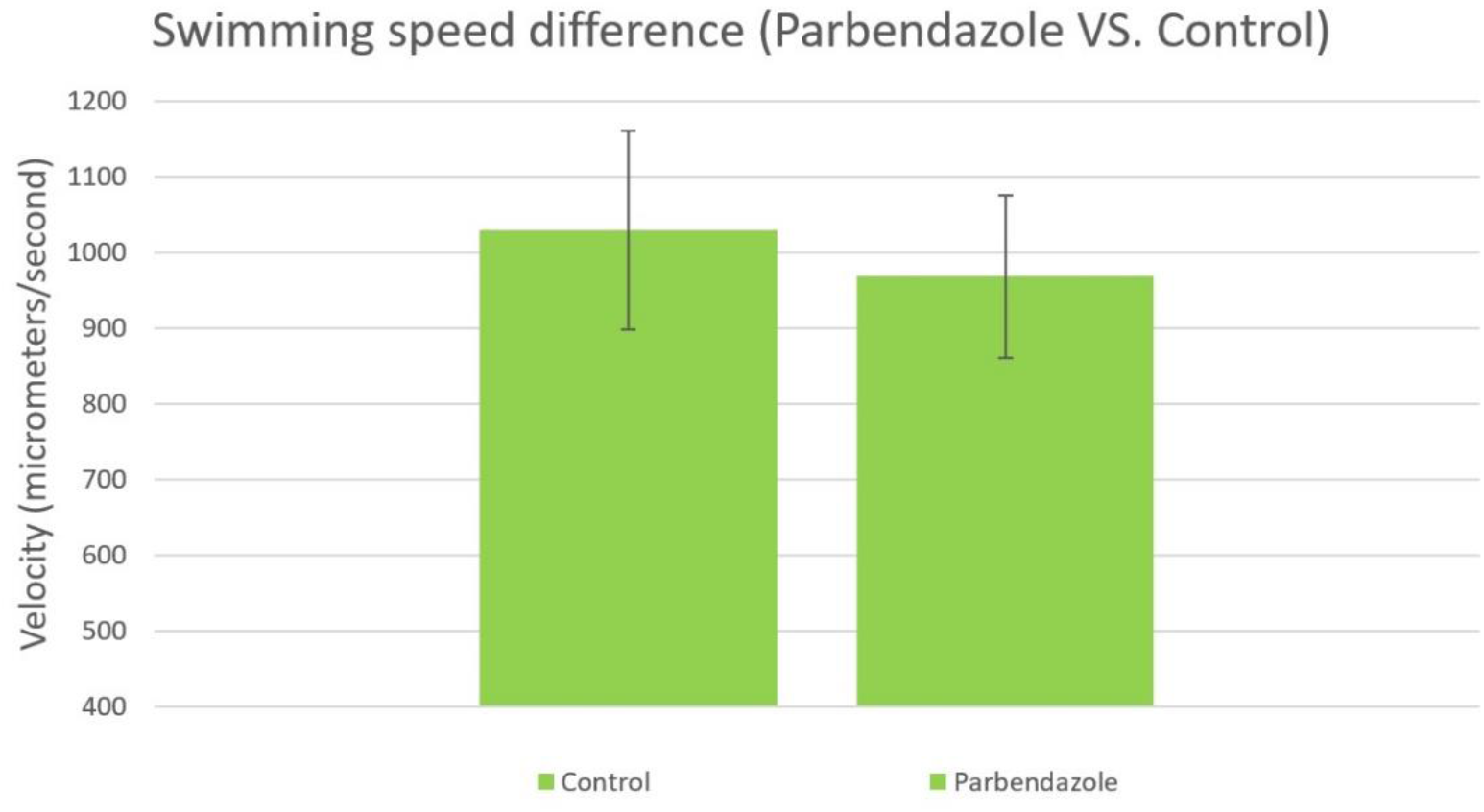
Swimming speed of paramecia without drug treatment(control) and 24 hours after parbendazole administration did not show a significant change (independent *t*-test, P>0.05, n=10). Error bars are ± SEM.

## III. DISCUSSION

### A. Implications for information processing and storage in microtubules

In the present study, we have shown that the disruption of microtubules will not cause significant impairment in learning behavior of P. caudatum. Therefore, microtubules may not be necessary for memory storage and learning in P. caudatum. It is noteworthy to mention that although parbendazole concentration was two orders of magnitude bigger than the IC50 concentration of parbendazole in Paramecium tetraurelia and it inhibited cellular growth in an effective fashion, it did not affect the learning behavior of P. caudatum. Meanwhile, it might be argued that parbendazole only affects dynamic microtubules and stable microtubules are untouched by parbendazole. However, it was shown that even stable microtubules reassemble after 6.5 hours (27) and parbendazole treatment for 24 hours can inhibit the reformation of stable microtubules. This has been supported by the fact that treatment of Vero cells for 21 hours with a 20 micromolar concentration of parbendazole causes total disassembly of microtubules(22). Since 24-hour treatment of cells with a concentration of 100 micromolar is well above what was reported in Vero cells, it is reasonable to assume that parbendazole has been able to affect both dynamic and stable microtubules. However, it is noteworthy to mention that in Havercroft et al. study(22), centrioles were not harmed by parbendazole in a variety of doses and durations (20 micromolar for 21 hours or 2 micromolar for 45 hours).

Accordingly, previous suggestions about the involvement of tubulins and microtubules in information storage in cells may need further revisitation. Additionally, if microtubules are not involved in information storage, it is hard to argue that they may be involved in information processing. Therefore, It might be reasonable to doubt the role of microtubules as fundamental structures which can support intelligent behavior(s) in an extensive range of species, as suggested by some scholars (5,28). However, since parbendazole does not affect centrioles, our results do not rule out the possibility of centrioles being involved in information processing as it was suggested by G Albrecht-Buehler (10–12).

### B. Learning in paramecium; Phototaxis, effects and aftereffects of electrical stimulation

Our results show that learning in P. caudatum can be impaired upon the administration of a GABA receptor partial allosteric modulator, however, not by disrupting the microtubular dynamics. We cannot explain exactly how this agent disrupts learning in paramecium, but perhaps it may be explained based on the previously proposed model (29), where learning in paramecium includes three main processes: a) light detection and phototaxis, b) disruption of phototaxis through electrical stimulation, and c) stimulation aftereffects.

### C. Light detection and phototaxis in paramecium

It is believed that phototaxis mechanisms in eukaryotes follow a straightforward rule (30). A photosensor molecule codes the light intensity and sends signals to a motor actuator for locomotion. In paramecium genus, Paramecium bursaria is a good example for phototaxis. Swimming speed of this ciliate decreases upon exposure to strong bright light and its membrane depolarizes in this process(31). This photosensitivity is independent from the symbiotic alga that exists in Paramecium bursaria as both chlorella-containing and chlorella free forms of the Paramecium bursaria show photosensitivity (31). On the other hand, its swimming speed increases upon normal light exposure due to hyperpolarization of its membrane potential (32). Interestingly, it is known that this membrane hyperpolarization causes an increased ciliary beat frequency(33) mediated through cyclic adenosine monophosphate (cAMP) molecules (32). While retinal was extracted from Paramecium bursaria as a possible chromophore molecule (34), the exact identity of photosensor molecule in P. caudatum is still unknown. It seems reasonable to assume that paramecium uses cAMP as an intermediate messenger to coordinate between its photosensor and cilia. We suggest that a similar molecular pathway can be responsible for phototaxis in P. caudatum. Since freely swimming paramecia spend only 39% of their time in the bright side of the trough (≈ 100 seconds out of 270, see figure 2), it can be suggested that P. caudatum shows a photophobic behavior similar to P. bursaria which involves cAMP, membrane potential fluctuations, and an unknown photosensor.

### D. Disruption of phototaxis through electrical stimulation

It is known that paramecium membrane contains Ca2+ channels (35) used in movement behaviors of the organism(33,36,37). Interestingly, it was shown that membrane depolarization can cause a reversal in ciliary beating direction of the organisms by changing the flow of Ca2+ ions across the membrane (38). Since the resting membrane potential of paramecium is around −25 millivolts (38), it is fair to assume that successive cathodal shocks will cause an outward flow of positive ions (and Ca2+ in particular) and cancellation of P. caudatum’s photophobic behavior by reducing its swimming speed (Fig. 6).

**FIG. 6.**
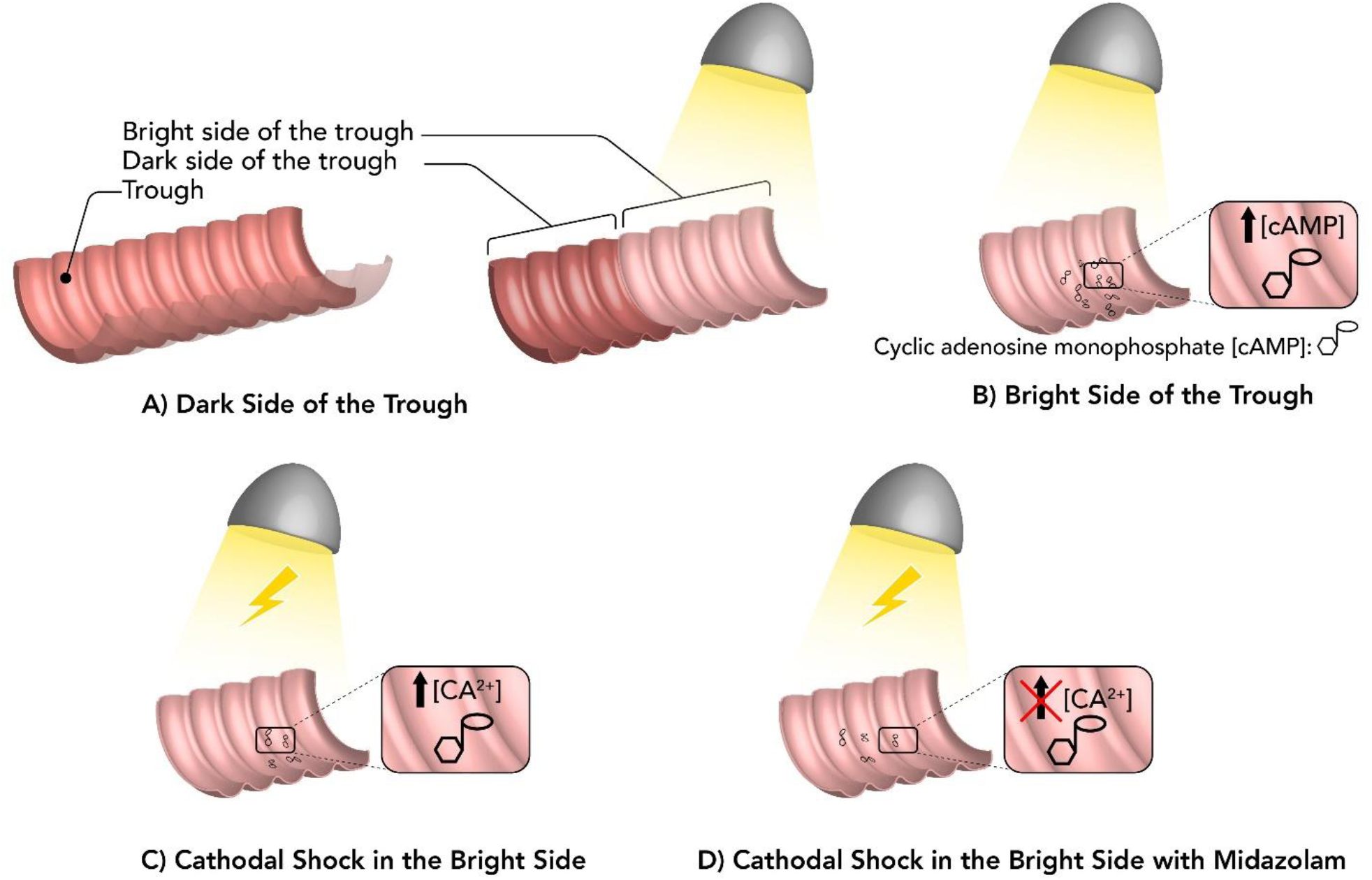
A schematic representation of the proposed model for learning in paramecium. A) When paramecium is swimming in the dark side of the trough, there is a baseline cAMP concentration which maintains a normal swimming speed. B) When paramecium enters the bright side of the trough, light exposure causes an increase in cAMP levels which consequently increase the swimming speed. C) When paramecium enters the bright side of the trough and receives successive electrical shocks, electrical shocks will cause subtle and temporary backward movements which leads to a normal swimming speed for paramecium almost equal to its swimming speed in the dark side. This causes accumulation of cAMP molecules in the cytosol which eventually cancels the photophobic behavior of paramecium during test trials. It seems to be a possible mechanism for learning in paramecium. D) Administration of midazolam as a GABA PAM will cancel the backward movements as mentioned in the text. This process can eliminate the effect of electrical shocks and the subsequent accumulation of cAMP molecules in paramecium; therefore, it will disrupt the learning behavior of paramecium.

### E. Stimulation aftereffects

Our assumptions to this point can explain why paramecia in the training group are relatively attracted to the bright side, but they do not explain the mechanisms of memory retention in paramecium. We believe that memory retention in this organism is related to the stimulation aftereffects. Light exposure produces a substantial amount of cAMP in the cytosol while the reduction of Ca2+ in the cell-due to electrical shocks-opposes the effect of cAMP as a speed booster. In the test trials, when there is no electrical shock, the stored cAMP exists in excessive amounts and it speeds up paramecium movement regardless of its position in the bright or dark side of the trough. This is in line with the experimental finding that paramecia in experimental group spend an almost equal amount of time in both halves of the trough (Fig. 2). We suggest that memory in P. caudatum in this task is encoded in the concentration of cAMP molecules.

### F. Pharmacological manipulations of learning in *P. caudatum*

Our results indicate that a benzodiazepine drug (a GABA receptor Partial Allosteric Modulator) can disrupt the learning behavior of P. caudatum. As shown in the previous reports, a GABAB agonist (baclofen) can cancel the reversal movement of Paramecium primaurelia and its effect can be blocked by a Ca2+ channel blocker (verapamil) (39). Midazolam as a GABA PAM opposes the effect of electrical shocks and helps the paramecium to exert its normal photophobic behavior towards the light (see Fig. 6 for more information).

### G. limitations

One of the main limitations of the current study is the lack of behavioral data for multiple doses of parbendazole or midazolam. Since our main goal in the present study was to identify (and not quantify) agents involved in paramecium learning, we only studied single non-toxic high concentrations of parbendazole and midazolam. However, lower dosages of the same pharmacologic agents might have different effects. Additionally, we used population density as a proxy for parbendazole’s effectiveness while a more direct measure (such as immunocytochemistry) might be a more straightforward technique to illustrate the effects of parbendazole on microtubular structure.

## V. CONCLUSION

In the present study, we have shown that microtubular networks in the cytosol do not seem to be major players in the learning process of P. caudatum. Instead, ionic flow disruption by a GABA receptor allosteric modulator can impair memory in P. caudatum. We disrupted microtubular dynamics in paramecium by using parbendazole to see its effect on paramecium’s learning process. We observed that a partial allosteric modulator of GABA (midazolam) could disrupt learning process in paramecium, but parbendazole could not. Our results suggest that microtubules are probably not involved in information storage in P. caudatum and there might be other mechanisms for this process. This finding may have significant consequences for theories that consider a major role for microtubules in information storage or processing in a wide range of animals including paramecium.

Further studies on the molecular cascade involving cyclic monophosphate and calcium ions could explain the learning behavior of P. caudatum. We suggested a molecular pathway to explain the learning behavior in P. caudatum based on the effect of a GABA PAM on this phenomenon, which can be further evaluated through future experiments. To this end, one experimental test can be using the calcium channel blockers. If reversal beating direction (i.e. due to depolarization and calcium influx) is the main mechanism responsible for P. caudatum learning, Ca2+ channel blockers should be able to neutralize learning behavior of P. caudatum.

## VI. ACKNOWLEDGEMENTS

This work has been supported by the shiraz university of medical sciences (Pharm.D. thesis grant). The authors would like to thank Dr. Vahid Salari for his thoughtful comments and Mehdi Aslani, the physics student at Isfahan University of Technology, for enhancing the quality of Figures 1 and 7.

## VII. AUTHOR CONTRIBUTIONS

AA and HS have proposed the idea, AA performed the experiments, AA, GH, and HS contributed to the development and completion of the idea, AA, GH, and HS analyzed the results, AA, GH, and HS participated in discussions and writing the manuscript.

## VIII. ADDITIONAL INFORMATION

The authors declare no competing interests. This study was funded by shiraz university of medical sciences as a part of Pharm.D. thesis of AA.

## References

1. Fröhlich H. Long Range Coherence and the Action of Enzymes. Nature. 1970 Dec;228(5276):1093.

2. Fröhlich H. The extraordinary dielectric properties of biological materials and the action of enzymes. PNAS. 1975 Nov 1;72(11):4211–5.

3. Hameroff S, Penrose R. Orchestrated reduction of quantum coherence in brain microtubules: A model for consciousness. Mathematics and Computers in Simulation. 1996 Apr 1;40(3):453–80.

4. Ekert A., Jozsa R., Penrose R., Stuart Hameroff. Quantum computation in brain microtubules? The Penrose–Hameroff ‘Orch OR’ model of consciousness. Philosophical Transactions of the Royal Society of London Series A: Mathematical, Physical and Engineering Sciences. 1998 Aug 15;356(1743):1869–96.

5. Koch C, Hepp K. Quantum mechanics in the brain. Nature. 2006 Mar;440(7084):611.

6. Beck F, Eccles JC. Quantum aspects of brain activity and the role of consciousness. PNAS. 1992 Dec 1;89(23):11357–61.

7. Schlosshauer MA. Decoherence: and the Quantum-To-Classical Transition [Internet]. Berlin Heidelberg: Springer-Verlag; 2007 [cited 2019 Jun 19]. (The Frontiers Collection). Available from: https://www.springer.com/gp/book/9783540357735

8. Salari V, Naeij H, Shafiee A. Quantum Interference and Selectivity through Biological Ion Channels. Scientific Reports. 2017 Jan 30;7:41625.

9. Salari V, Moradi N, Sajadi M, Fazileh F, Shahbazi F. Quantum decoherence time scales for ionic superposition states in ion channels. Phys Rev E. 2015 Mar 9;91(3):032704.

10. Albrecht-Buehler G. Cellular infrared detector appears to be contained in the centrosome. Cell Motility. 1994;27(3):262–71.

11. Albrecht-Buehler G. Changes of cell behavior by near-infrared signals. Cell Motility. 1995;32(4):299–304.

12. Albrecht-Buehler G. Autofluorescence of Live Purple Bacteria in the Near Infrared. Experimental Cell Research. 1997 Oct 10;236(1):43–50.

13. Hameroff S, Penrose R. Consciousness in the universe: A review of the ‘Orch OR’ theory. Physics of Life Reviews. 2014 Mar 1;11(1):39–78.

14. Craddock TJA, Tuszynski JA, Hameroff S. Cytoskeletal Signaling: Is Memory Encoded in Microtubule Lattices by CaMKII Phosphorylation? PLOS Computational Biology. 2012 Mar 8;8(3):e1002421.

15. Hameroff SR, Craddock TJA, Tuszynski JA. “MEMORY BYTES” — MOLECULAR MATCH FOR CaMKII PHOSPHORYLATION ENCODING OF MICROTUBULE LATTICES. J Integr Neurosci. 2010 Sep 1;09(03):253–67.

16. Armus HL, Montgomery AR, Gurney RL. Discrimination Learning and Extinction in Paramecia (P. Caudatum). Psychol Rep. 2006 Jun 1;98(3):705–11.

17. Alipour A, Dorvash M, Hatam G, Yeganeh Y. Paramecium Learning: New Insights. J Protozool Res. 2018;28(1–2):22–32.

18. Zhou M, Xia H, Xu Y, Xin N, Liu J, Zhang S. Anesthetic action of volatile anesthetics by using Paramecium as a model. J Huazhong Univ Sci Technol [Med Sci]. 2012 Jun 1;32(3):410–4.

19. Armus HL, Montgomery AR, Jellison JL. DISCRIMINATION LEARNING IN PARAMECIA (P. caudatum). The Psychological Record; Heidelberg. 2006 Fall;56(4):489–98.

20. Al-Hadiya BMH. Chapter 5 - Parbendazole. In: Brittain HG, editor. Profiles of Drug Substances, Excipients and Related Methodology [Internet]. Academic Press; 2010 [cited 2019 Jun 19]. p. 263–84. Available from: http://www.sciencedirect.com/science/article/pii/S1871512510350059

21. Davidse LC. Benzimidazole Fungicides: Mechanism of Action and Biological Impact. Annual Review of Phytopathology. 1986;24(1):43–65.

22. Havercroft JC, Quinlan RA, Gull K. Binding of parbendazole to tubulin and its influence on microtubules in tissue-culture cells as revealed by immunofluorescence microscopy. Journal of Cell Science. 1981 Jun 1;49(1):195–204.

23. Pape R, Kissmehl R, Glas-Albrecht R, Plattner H. Effects of anti-microtubule agents on Paramecium cell culture growth. European Journal of Protistology. 1991 Sep 9;27(3):283–9.

24. Reves JG, Fragen RJ, Vinik HR, Greenblatt DJ. Midazolam: pharmacology and uses. Anesthesiology. 1985 Mar;62(3):310–24.

25. Olkkola KT, Ahonen J. Midazolam and Other Benzodiazepines. In: Schüttler J, Schwilden H, editors. Modern Anesthetics [Internet]. Berlin, Heidelberg: Springer Berlin Heidelberg; 2008 [cited 2019 Jun 19]. p. 335–60. (Handbook of Experimental Pharmacology). Available from: https://doi.org/10.1007/978-3-540-74806-9_16

26. Crevat-Pisano P, Dragna S, Granthil C, Coassolo P, Cano JP, Francois G. Plasma concentrations and pharmacokinetics of midazolam during anaesthesia. Journal of Pharmacy and Pharmacology. 1986;38(8):578–82.

27. Schulze E, Kirschner M. Dynamic and stable populations of microtubules in cells. The Journal of Cell Biology. 1987 Feb 1;104(2):277–88.

28. Litt A, Eliasmith C, Kroon FW, Weinstein S, Thagard P. Is the Brain a Quantum Computer? Cognitive Science. 2006 May 6;30(3):593–603.

29. Alipour A, Dorvash M, Yeganeh Y, Hatam G, Seradj SH. Possible Molecular Mechanisms for Paramecium Learning. Journal of Advanced Medical Sciences and Applied Technologies. 2017;3(1):39–46.

30. Jékely Gáspár. Evolution of phototaxis. Philosophical Transactions of the Royal Society B: Biological Sciences. 2009 Oct 12;364(1531):2795–808.

31. Matsuoka K, Nakaoka Y. Photoreceptor Potential Causing Phototaxis of Paramecium Bursaria. Journal of Experimental Biology. 1988 Jul 1;137(1):477–85.

32. Mitarai A, Nakaoka Y. Photosensitive Signal Transduction Induces Membrane Hyperpolarization in Paramecium bursaria. Photochemistry and Photobiology. 2005;81(6):1424–9.

33. Machemer H. Frequency and directional responses of cilia to membrane potential changes inParamecium. J Comp Physiol. 1974 Sep 1;92(3):293–316.

34. Tokioka R, Matsuoka K, Nakaoka Y, Kito Y. EXTRACTION OF RETINAL FROM Paramecium bursaria. Photochemistry and Photobiology. 1991;53(1):149–51.

35. Thiele J, Schultz JE. Ciliary membrane vesicles of paramecium contain the voltage-sensitive calcium channel. PNAS. 1981 Jun 1;78(6):3688–91.

36. Doughty MJ, Dryl S. Control of ciliary activity in Paramecium: An analysis of chemosensory transduction in a eukaryotic unicellular organism. Progress in Neurobiology. 1981 Jan 1;16(1):1–115.

37. Hinrichsen RD, Saimi Y, Hennessey T, Kung C. Mutants in paramecium tetraurelia defective in their axonemal response to calcium. Cell Motility. 1984;4(4):283–95.

38. Nakaoka Y, Imaji T, Hara M, Hashimoto N. Spontaneous fluctuation of the resting membrane potential in Paramecium: amplification caused by intracellular Ca2+. Journal of Experimental Biology. 2009 Jan 15;212(2):270–6.

39. Bucci G, Ramoino P, Diaspro A, Usai C. A role for GABAA receptors in the modulation of Paramecium swimming behavior. Neuroscience Letters. 2005 Oct 7;386(3):179–83.

